# Modulation of hippocampal synaptic transmission by mast cells at the hippocampal-thalamic border during early brain development

**DOI:** 10.1101/2025.09.30.677297

**Authors:** Helena Gimeno Agud, Luke William Paine, Maartje Hof, Tommas Ellender

## Abstract

Synaptic development is a highly structured process and can be modulated by the biogenic amine, histamine, whose dysregulation during brain development can be part of the aetiology of the neurodevelopmental disorders Tourette’s syndrome and OCD. Well documented in the literature is the extensive histaminergic axonal innervation originating from neurons in tuberomammillary nucleus of the hypothalamus, however, less is known about non-neuronal sources of histamine, such as mast cells, and their potential functional importance in development. Here we set out to investigate the spatiotemporal localisation of mast cells in the developing C57Bl/6 mouse brain and the potential impact of their degranulation on developing synapses. Firstly, we find that the mouse brain contains relatively larger numbers of mast cells during the first and second postnatal week. Secondly, that in this period the highest density of mast cells is found just below the hippocampus at the hippocampal-thalamic border with lower numbers found in other brain regions. Thirdly, using the mast cell degranulator C48/80 we observe that their degranulation can facilitate synaptic transmission at the perforant pathway inputs to the ventral portion of the dentate gyrus which could not be blocked by co-application of the histamine H_3_ receptor antagonist thioperamide. In contrast histamine superfusion tended to decrease synaptic transmission at this pathway which was sensitive to thioperamide co-application. In conclusion, these results suggest that mast cells are present in the developing C57Bl/6 mouse brain and in particular at the hippocampal-thalamic border where upon degranulation they appear to modulate hippocampal synaptic transmission.

## Introduction

The neuromodulator histamine regulates key neurophysiological processes in both the developing and mature nervous system including neurogenesis (Molina-Hernández & Velasco, 2008), synaptic transmission (Ellender *et al*., 2011; Márquez-Gómez *et al*., 2023) and synaptic plasticity (Brown *et al*., 1995; Han *et al*., 2020) amongst others (Schwartz *et al*., 1991; Haas & Panula, 2003; Takahashi *et al*., 2006; Panula *et al*., 2014; Han *et al*., 2020; Lucaci *et al*., 2023). Histaminergic signalling has also been shown to be dysregulated in various neurological and neurodevelopmental disorders (Ercan-Sencicek, 2010; Fernandez *et al*., 2012; Karagiannidis *et al*., 2013; Baldan *et al*., 2014; Shan *et al*., 2015; Robertson *et al*., 2017; Carthy & Ellender, 2021; Xu *et al*., 2022; Ma *et al*., 2023). The early onset of these disorders suggest that histamine likely impacts neuronal and circuit development and this has led to an increased interest both in understanding the physiological and pathological roles for histamine during early developmental stages (Panula *et al*., 2014; Rapanelli *et al*., 2017; Han *et al*., 2020; Valle-Bautista *et al*., 2021) as well as its potential as a therapeutic target (Rapanelli & Pittenger, 2016; Zhang et al., 2020; Carthy & Ellender, 2021). Histamine is detectable early in brain development and likely comes from diverse sources, and as postnatal development progresses the main source shifts to the neurons of the hypothalamic tuberomammillary nucleus (TMN) (Vanhala *et al*., 1994; Molina-Hernández *et al*., 2012) which extend axons and progressively innervate much of the central nervous system (CNS) (Haas & Panula, 2003; Haas et al., 2008; Lin et al., 2023). Although histamine-expressing neurons in the TMN are seen from the first postnatal week onwards in rodents (Reiner, 1988; Panula et al., 2014), many regions of CNS appear to not contain detectable histaminergic afferents until much later in development and for some regions, including the striatum, they remain consistently sparse (Han *et al*., 2020; Lin *et al*., 2023; Márquez-Gómez *et al*., 2023). This raises the question, what other sources of histamine might exist that could potentially impact developing neurons and circuits in these brain regions.

Mast cells (MCs) are bone-marrow-derived immune cells known for their capacity to synthesize and release a plethora of preformed mediators, including histamine, by which they control various inflammatory and allergic processes in the body (Baldwin, 2006; St John *et al*., 2023). They are also found in the peripheral nervous system (Van Nassauw *et al*., 2007) and the CNS (Silver *et al*., 1996; Silver & Curley, 2013) where they have been suggested to contribute significant amounts of histamine (Ferrer *et al*., 1979; Yamatodani *et al*., 1982) and are found at high numbers in specific regions at distinct stages of life (Auvinen & Panula, 1988; Panula *et al*., 1988; Lambracht-Hall *et al*., 1990a; Khalil *et al*., 2007; Lenz *et al*., 2018) where they can modulate local neurons and immune cells (Theoharides *et al*., 1995; Kovacs *et al*., 2006; Dong *et al*., 2017; Lenz *et al*., 2018; Flores *et al*., 2019; Forsythe, 2019; Theoharides *et al*., 2024; Blanchard *et al*., 2025). Although much focus has been on mast cells in rats (Auvinen & Panula, 1988; Panula *et al*., 1988; Lambracht-Hall *et al*., 1990a; Khalil *et al*., 2007; Blanchard *et al*., 2025) far less is known about their distribution in mice and whether the histamine they contain is capable of modulating physiological processes, including synaptic transmission.

Here we assessed the presence of mast cells in several key CNS regions in C57Bl6/J mice, focussing on the cortex, striatum, thalamus, hypothalamus and hippocampus, during early development using various immunocytochemical approaches. We then explored whether pharmacologically induced degranulation of these mast cells could impact synaptic transmission.

## Materials and Methods

### Animals and housing

All experiments were conducted on C57Bl/6 wildtype or the B6.Cg-Kit<W-sh>/HnihrJaeBsmJ transgenic mice which has been bred on a C57Bl/6 background (https://www.jax.org/strain/030764) and exhibits depleted mast cells numbers. Mice were used of both sexes with *ad libitum* access to food and water. All mice were bred and housed in a temperature-controlled animal facility (normal 12:12 h light/dark cycles) and used in accordance with the UK Animals (Scientific Procedures) Act (1986) and European Ethics Committee (decree 2010/63/EU) and approved by the Committee on Animal Care and Use at the University of Oxford, UK and the Ethische Commissie Dierproeven at the University of Antwerp, Belgium.

### Histological Analyses

The brains of mice were immersed and fixed for 3 days in cold 4% paraformaldehyde (PFA) in 0.1 M PBS and subsequently kept in PBS until used. Coronal 50 µm sections were made on a vibratome (Leica 1000S) and washed (PBS, 3x, each 10 min.). Toluidine blue (TB) staining was performed by incubating sections for 10 sec. in a solution of 4 mg/ml TB in 60% Ethanol at pH 2.0 (Nautiyal *et al*., 2012; Dong *et al*., 2017). Slices then underwent a series of ethanol dehydration steps: 50% ethanol for 15s, 70% 45s, 95% 1 min (x2), 100% 1 min (x2). Slices were cleared in xylene for 2 mins (x3). Finally, slides were cover slipped with Eukitt mounting medium (Sigma-Aldrich Catalog #05393). TB-labelled mast cells in brain tissue were identified based on the colour, shape and size of cells. TB stains MCs red/purple with the background tissue stained blue. For immunohistochemical staining the sections were washed in PBS-Tx (0,3%) three times before adding the primary antibody in solution with PBS-Tx (0,3%). Slices with 1:500 avidin-FITC (Merck-Millipore, 189727-1ml) or 1:2000 avidin-Sulforhodamine101 (sigma, A2348-5MG) antibodies were incubated for 2h at room temperature under agitation, after which they were washed in PBS and mounted. Slices with 1:200 anti-rat mcpt6 (R&D systems, MAB3736) and 1:50 anti-rat c-kit (Bio-Techne Sales Corp, AF13556) antibodies were incubated overnight at 4°C under agitation. After this, they were washed with PBS for three times. Secondary antibody, FITC for c-kit and Cy3 for mcpt6, were added in solution with PBS-Tx (0,3%) and incubated for 2h at room temperature under agitation. Finally, all slices were mounted in Vectashield + DAPI and visualised with an epifluorescence microscope. Images were taken with an Olympus Microscope equipped with a Hamamatsu or ThorCam camera and acquired with HCImage Live software and processed with ImageJ.

### Electrophysiology

Acute striatal slices were made from mice between postnatal day (P)7–11 of age. Mice were anesthetized with isoflurane and rapidly decapitated. Coronal 400 µm slices were cut using a vibrating microtome (Microm HM650V). Slices were prepared in aCSF containing the following (in mm): 65 sucrose, 85 NaCl, 2.5 KCl, 1.25 NaH_2_PO_4_, 7 MgCl_2_, 0.5 CaCl_2_, 25 NaHCO_3_, and 10 glucose, pH 7.2–7.4, bubbled with carbogen gas (95% O_2_/5% CO_2_). Slices were immediately transferred to a storage chamber containing aCSF (in mM) as follows: 130 NaCl, 3.5 KCl, 1.2 NaH_2_PO_4_, 2 MgCl_2_, 2 CaCl_2_, 24 NaHCO_3_, and 10 glucose, pH 7.2–7.4, at 32°C and bubbled with carbogen gas until used for recording. Within the interface chamber, slices were continuously superfused with recording aCSF (see above) and bubbled with carbogen gas. Borosilicate glass recording electrodes were filled with aCSF and placed in either the ventral dentate gyrus (DG) or stratum radiatum of the CA1 region of the hippocampus adjacent to a bipolar stimulating electrode. Recordings were made using a MultiClamp 700A amplifier (Axon Instruments, Molecular Devices, CA, USA), a NI-DAQ and WINWCP software. Hippocampal afferents were stimulated every 10 sec. at approximately half-maximal fEPSP (field excitatory postsynaptic potential) amplitude using a DS2 isolated stimulator. For each slice, a 10-20 min. baseline period was recorded to ensure sufficient stability of the fEPSP amplitude. After a baseline recording, the tissue was superfused with recording aCSF containing the mast cell degranulator compound 48/80 (C48/80, 1 µg/ml) and fEPSPs were recorded for 20 min. and finally by a further 20 min. of superfusion with C48/80 and histamine (10 µM). For data analysis, custom-written scripts were utilised in Igor Pro (Wavemetrics RRID:SCR_000325). The fEPSP traces included a stimulation artefact, first negative deflection (fibre volley indicating a pre-synaptic action potential), and second negative deflection (indicative of synaptic transmission). From the second deflection the peak amplitude (the difference between baseline voltage and fEPSP peak) was obtained.

### Statistical analysis and data representation

All data are presented as means ± SEM. Statistical tests were all two-tailed and performed using SPSS 17.0 (IBM SPSS statistics, RRID:SCR_002865). Continuous data were assessed for normality and appropriate parametric (ANOVA, paired *t*-test and unpaired *t*-test) or non-parametric (Mann-Whitney) statistical tests were applied (* p<0.05, ** p<0.01, *** p<0.001).

## Results

### Assessing toluidine blue labelled cells in the developing mouse brain

Firstly, the number of mast cells in the C57Bl/6 mouse brain were counted at various developmental stages ranging from embryonic day (E)17.5 to postnatal day (P)28 and older. Mast cells (MCs) were visualised in 50 µm coronal sections of paraformaldehyde-fixed mouse brains using the metachromatic dye TB, which stains mast cells a purplish colour against a blue background. Their numbers were then assessed across various regions (**Figure 1A, B**). The focus was on the striatum as this region exhibits very sparse histaminergic innervation (Han *et al*., 2020; Márquez-Gómez *et al*., 2023) as well as the hippocampus, thalamus and hypothalamus as some of these regions had previously been shown to contain high numbers of mast cells in other species. These regions were examined in their entirety for TB purplish cells of approximately 20 μm in diameter and containing granules and located outside of blood vessels; cells meeting these criteria were counted as mast cells. Counts were made in at least 9 sections per mouse brain spanning the start of rostral cortical regions M1/S1 (Paxinos Figure 19) including the striatum and ending at the ventral hippocampal regions (Paxinos Figure 50). We find that overall mast cell numbers increase rapidly after birth and are found in large numbers during the initial postnatal weeks as can be seen when their absolute numbers are averaged across all counted sections for each brain (E17.5: 0.24 ± 0.08, P3-6: 8.82 ± 2.34, P9-12: 3.03 ± 1.07, P28+: 1.38 ± 0.69, n = 8, 8, 8 and 4 brains, **Figure 1C**). This reveals that the highest numbers of toluidine labelled cells are found during the first postnatal week (P3-6 vs E17.5; p=0.0007 and P3-6 vs P9-12; p=0.039, ANOVA, **Figure 1C**). We next characterised the location of mast cells within specific brain regions at this developmental stage. Regions were assessed conservatively and in the majority of sections we were able to confidently determine the location of mast cells with the help of a mouse brain atlas (P3-6: 84% and P9-12: 95% of all cells) (Franklin, 2007). Normalising over all mast cells counted within these regions consistently revealed that the hippocampus exhibited a large number of mast cells as compared to other brain regions both in the first and second postnatal week (P3-6: 61.78 ± 14.86% and P9-12: 66.60 ± 8.81%, p=0.0006 and p=3.3E-6, ANOVA, both n=8 brains, **Figure 1D**). No statistically significant differences were found between the thalamus, hypothalamus and striatum in the proportion of MCs and all contained less than 10% of MCs. The MCs in the cortex were significantly more abundant than in the thalamus and hypothalamus (cortex vs thalamus; p=0.009 and cortex vs hypothalamus; p=0.032, Kruskal-Wallis test, both n=8 brains). In conclusion, these results suggest that MCs are not seen in large numbers in the striatum, which exhibits sparse histaminergic innervation, but are present in higher numbers in hippocampal regions.

**Figure 1.**
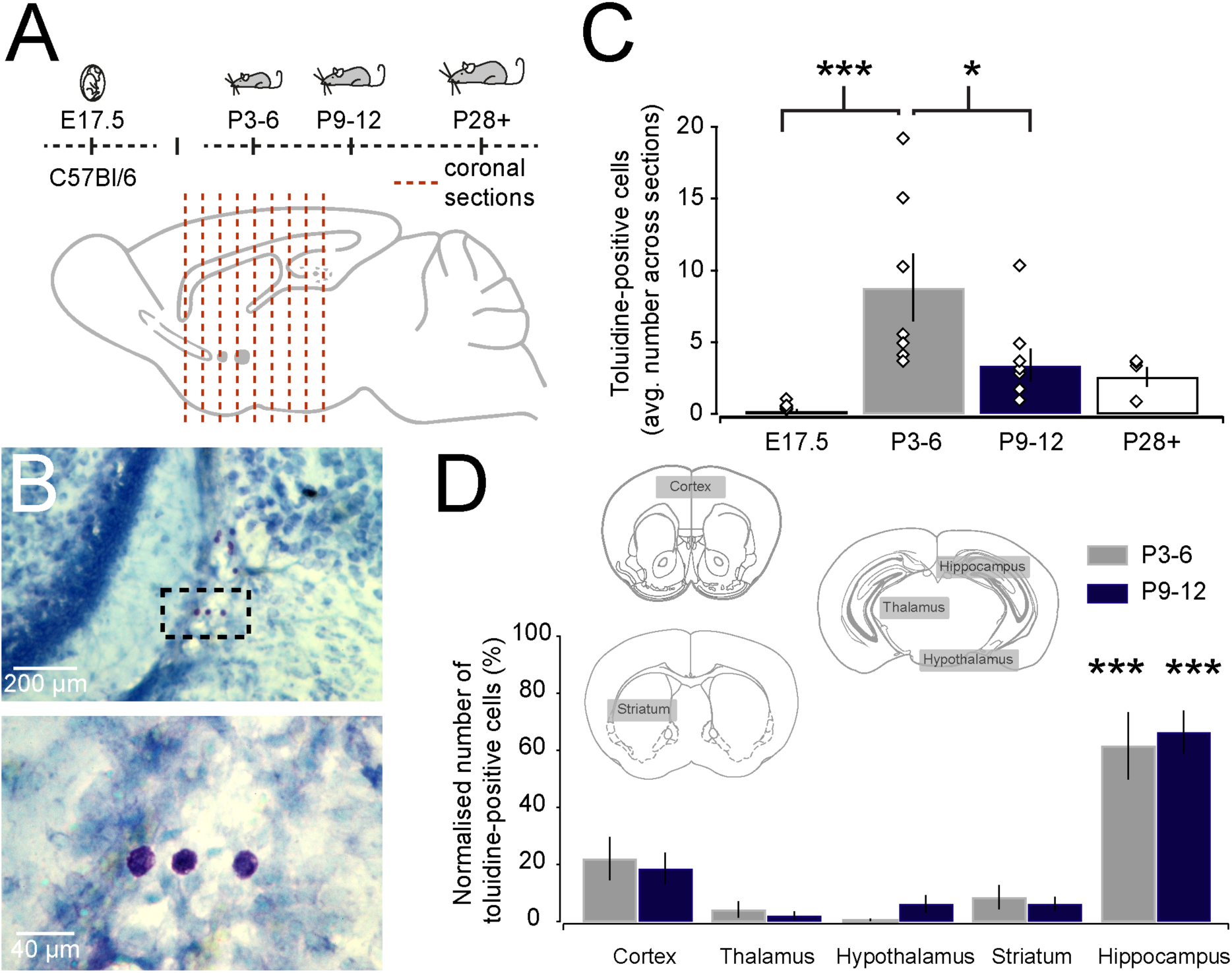
Mast cell counts in the developing mouse brain using toluidine blue staining. (**A**) Mouse brain sections ranging from the most rostral portion of striatum (Add Paxinos Figure 19) up to caudal hippocampus (Paxinos Figure 50) at different developmental stages were analysed for toluidine labelled cells. (**B**) Toluidine labelled cells were observed as darkly purple labelled cells. (**C**) Average mast cell numbers across all sections counted showed a peak in their numbers during the first postnatal week. (**D**) Barplot of normalised numbers of mast cells in specific brain regions revealed that the hippocampus contains the largest numbers as compared to other brain regions in both the first and second postnatal weeks. All data presented as mean ± SEM, * p<0.05; *** p<0.001.

### Assessing mast cell numbers in the hippocampal region

We next investigated in more detail the hippocampal distribution of these toluidine blue labelled cells. To do this we defined four distinct hippocampal regions; namely the ventral border between hippocampus and thalamus, the lateral fimbria region, the hippocampus proper including DG, CA3 and CA1 regions and the medial ventricular region (**Figure 2A**). Re-counting of the numbers of toluidine blue labelled cells within these distinct hippocampal regions revealed that significantly more are found at the ventral border between hippocampus and thalamus (79.39 ± 6.34%) as compared to other regions, and this was seen during both the first and second postnatal week (p = 0.0009 and p = 0.00001, ANOVA, both n = 8 brains, **Figure 2B**). Secondly, a separate set of brain sections were labelled with the glycoprotein avidin conjugated with FITC (avidin-FITC), which has been shown to also reliably label mast cells (Bergstresser *et al*., 1984; Tharp *et al*., 1985). Counting of avidin-FITC labelled cells in the various hippocampal regions in these sections (using similar criteria of their size, location and presence of granules) confirmed the high numbers of mast cells found at the border of hippocampus and thalamus during the first two postnatal weeks (P3-6: 64.19 ± 4.16% and P9-12: 68.11 ± 8.69%, p = 0.004 and p = 0.0008, ANOVA, both n = 4 brains, **Figure 2C**) which was analogous to the observations made using toluidine blue staining (P3-6 toluidine vs avidin FITC and P9-12 toluidine vs avidin FITC: both p > 0.05, Mann-Whitney, n = 8 and 4 brains). In addition, we explored whether this population of mast cells at the border of the hippocampus and thalamus could also be labelled with other known mast cell markers and whether their characteristics were similar. We found that indeed they could be labelled also with avidin conjugated to sulforhodamine101 (Kafitz *et al*., 2008; Joulia *et al*., 2015) as well as with an anti-mcpt6 antibody that detects mast cell protease-6 stored in mast cell granules (Abraham *et al*., 2007), and the c-kit or CD117 tyrosine kinase receptor, which is important for mast cell maturation (Tsai *et al*., 2022) (**Figure 2D**). We also looked in hippocampal sections of Mrgprb2-cre mice (McNeil *et al*., 2015) crossed with the ROSA26-tdTomato reporter mouse line and observed fluorescent cells at the hippocampal thalamic border (**Figure 2D**). Lastly, we stained brain sections from ckit (Kit<W-sh> transgenic (see Methods), which should contain overall significantly reduced numbers of mast cells at this age (Lyon & Glenister, 1982), with avidin-FITC and did not observe labelled cells within the hippocampal region that adhered to all our criteria (n = 4 brains tested, **Figure 2E**) although we did see some avidin-FITC labelled cells within blood vessels and the choroid plexus (**Figure 2E**) (Kitamura *et al*., 1978; Yamazaki *et al*., 1994; Grimbaldeston *et al*., 2005; Mukai *et al*., 2017; Kierdorf *et al*., 2019; Quintana, 2019). In conclusion, these data suggest that there is a transient population of mast cells in the hippocampus at the hippocampal-thalamic border during the first weeks of postnatal life.

**Figure 2.**
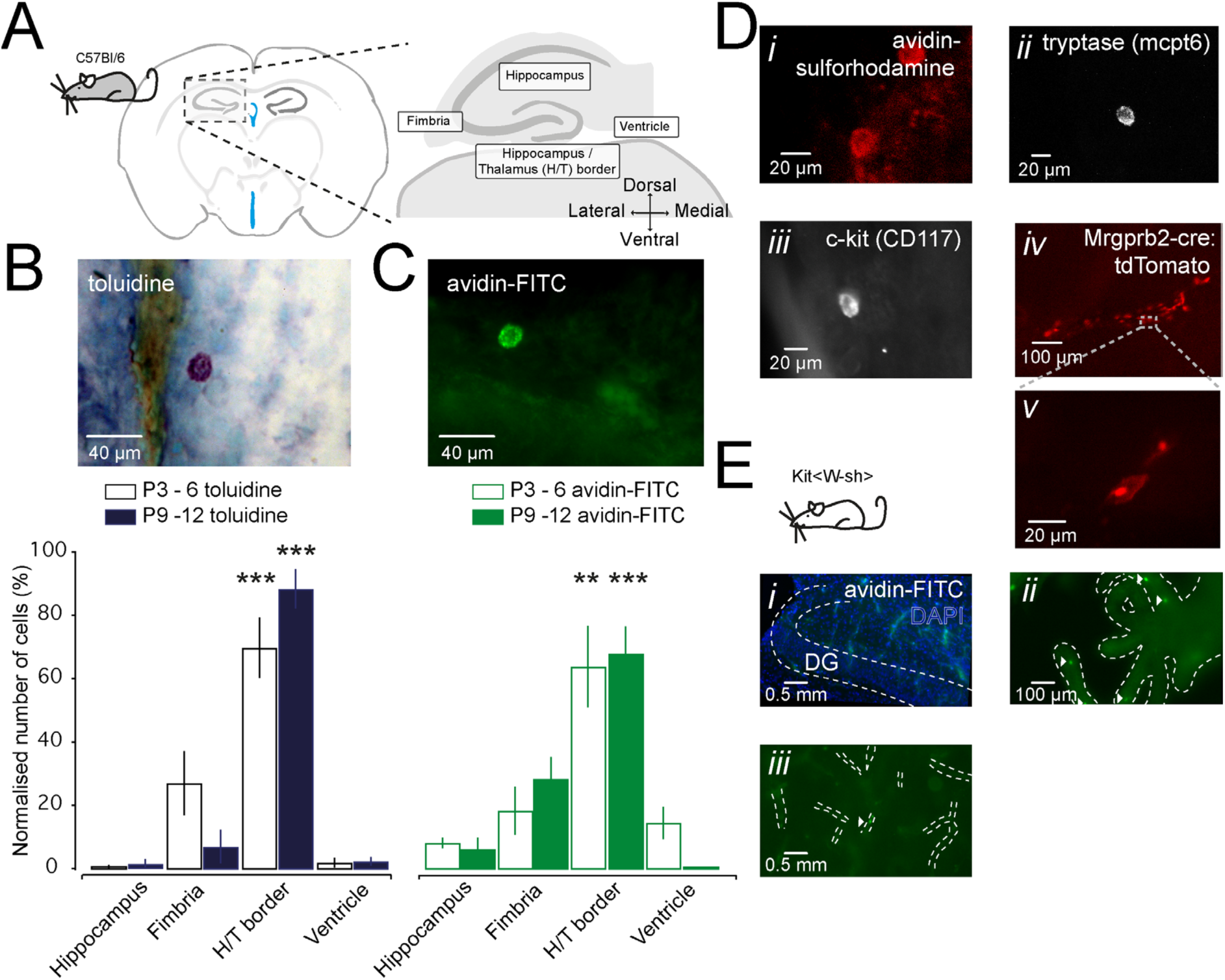
The ventral hippocampal-thalamic border is enriched for mast cells during the first and second postnatal week. (**A**) Analysis was focussed on 4 specific regions in the hippocampus, namely the ventral border between hippocampus and thalamus, the fimbria region, the hippocampal proper including the dentate gyrus, CA3 and CA1 regions and the ventricular region. (**B**) Most toluidine positive cells were found at the ventral border between hippocampus and thalamus with lower numbers found in other regions including the fimbria region (17.19 ± 6.52%), hippocampus (1.06 ± 0.79%) and the ventricular region (2.36 ± 0.93%). (**C**) Avidin-FITC staining further confirmed this finding with the greatest number of positive cells seen at the ventral hippocampal-thalamic border. (**D**) Other markers also revealed labelled cells in sections of the hippocampal-thalamic border with similar morphology **i** avidin-sulfurhodamine, **ii** tryptase (mcpt6 antibody), **iii** c-kit (CD117 antibody) **iv** Mrgprb2-cre:tdTomato mast cell-specific transgenic mice at low magnification (10x) and **v** higher magnification (40x). (**E**) No avidin-FITC labelled cells were seen in **i** sections containing hippocampus from ckit (Kit<W-sh> transgenic mice, but some labelled cells (white arrowheads) were seen in **ii** choroid plexus and **iii** within some blood vessels. All data presented as mean ± SEM, ** p<0.01; *** p<0.001.

### Assessing effect of degranulation of mast cells on synaptic transmission in the hippocampus

We next assessed whether degranulation of this transient population of mast cells and release of their content could modulate physiological processes in the hippocampus (**Figure 3**). We focussed on synaptic transmission at the perforant path in the ventral portion of the dentate gyrus as this region is proximal to the observed mast cells. Acute 400 μm thick hippocampal sections of P7-11 old C57Bl6/J mice were made and placed in an Oslo-style interface chamber. A tungsten stimulation electrode was positioned below the granule cell layer of the dentate gyrus to stimulate the perforant path afferents coming from the entorhinal cortex. Field recordings were made using a glass electrode placed more medial and downstream from the stimulating electrode below the granule cell layer (**Figure 3A**). A stimulation pulse was given every 10 seconds and this led to a reliable and stable field fEPSP as recorded with the glass electrode (**Figure 3A**). After a 10-minute baseline recording, the mast cell de-granulator C48/80 (1 μg/ml) (Paton, 1951) was superfused for 20 minutes and the effect on the fEPSP amplitude was assessed. We find that this led to a significant increase in the fEPSP amplitude (146.3 ± 24.8%, p=0.0011, paired t-test, n = 12, **Figure 3A**). We subsequently continued the superfusion of C48/80 but now also included histamine (10 μM) and found that this led to a trend in reducing the fEPSP amplitude (C48/80: 146.3 ± 24.8% and C48/80+histamine: to 123.7 ± 30.1%, p=0.23, paired t-test, n = 12, **Figure 3A**). To investigate whether changes in the fEPSP amplitude were mediated through action at histamine receptors we superfused histamine together with the H_3_ receptor antagonist thioperamide, as histamine had previously been shown to regulate hippocampal synaptic transmission through H_3_ histamine receptors in adult mice (Bekkers, 1993; Brown *et al*., 1995).The potent and selective H_3_ receptor antagonist thioperamide acts in the nM range in cell-based assays (Arrang *et al*., 1987; Hew *et al*., 1990; Arrang *et al*., 1995; Morisset *et al*., 2000; Molina-Hernandez *et al*., 2001; Gbahou *et al*., 2006) and in the μM range in brain slices studies (Arias-Montano *et al*., 2001) and for these reasons a moderately high concentration of thioperamide (10 μM) was used to guarantee a sufficient concentration at slices under these recording conditions. The presence of thioperamide did not impact the C48/80-mediated increase in amplitude (C48/80+thioperamide: to 172.5 ± 14.5%, p=0.0055, paired t-test, n = 5, **Figure 3A**) but exhibited a trend towards attenuating the negative modulation seen with histamine (C48/80+histamine: 123.7 ± 30.1%, and C48/80+histamine+thioperamide: 163.5 ± 25.9%, p=0.079, Mann-Whitney, n = 12 and n = 5, **Figure 3A**). The increase in the fEPSP amplitude upon superfusion of C48/80 was not seen when the experiment was performed in hippocampal slices made from Kit<W-sh> transgenic mouse which are depleted of brain mast cells (C48/80 to 108.0 ± 7.5%, p>0.05, paired t-test, n = 13, **Figure 3B**) but the subsequent addition of histamine (10 μM) did produce a trend towards a decrease in fEPSP amplitude (C48/80+histamine to 85.2 ± 12.4%, p=0.096, paired t-test, n = 13, **Figure 3B**). Lastly, to explore the spatial extent of the effect of C48/80 degranulation we recorded the impact on transmission at more distal synapses and specifically the Schaffer collateral synapses in the hippocampal CA1 region in slices of P7-11 old C57Bl6/J mice. This revealed that superfusion of C48/80 here did not affect synaptic transmission (C48/80+histamine to 97.3 ± 4.2%, p=0.14, Wilcoxon, n = 5, **Figure 3C**) but that histamine exhibited a trend towards decreasing the fEPSP amplitude (C48/80+histamine to 82.5 ± 6.3%, p=0.27, Wilcoxon, n = 4, **Figure 3C**). In conclusion, these results suggest that degranulation of mast cells can modulate synaptic transmission in close-by regions such as at the ventral dentate gyrus perforant pathway during early postnatal brain development and this might occur through a histamine-independent mechanism.

**Figure 3.**
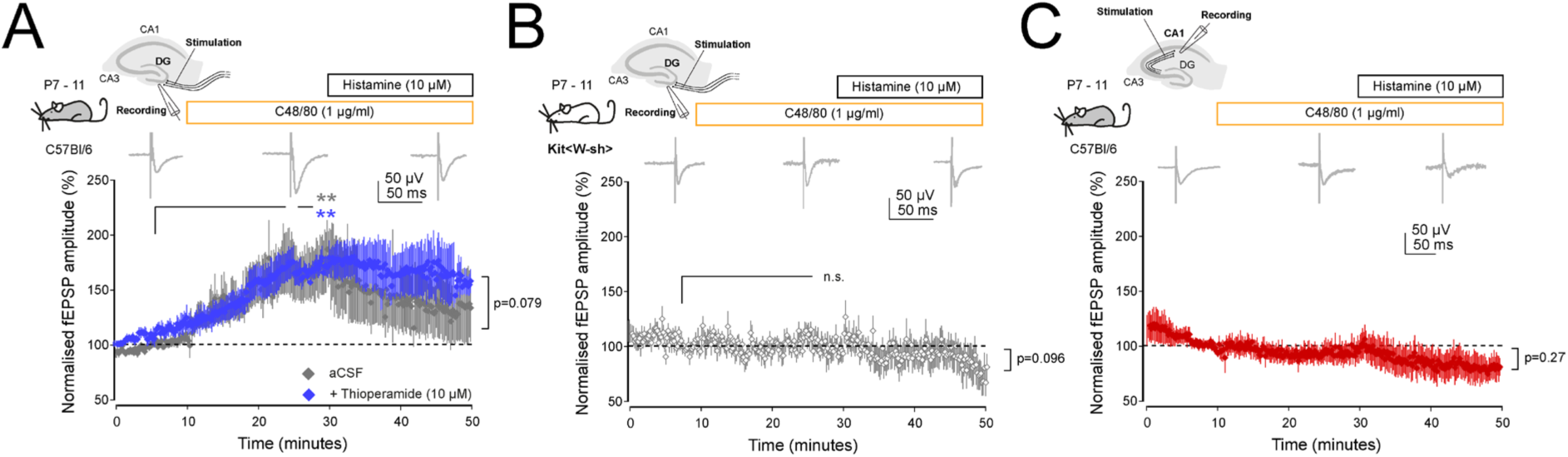
Impact of mast cell degranulation on hippocampal synaptic transmission. (**A**) Graph of the normalised fEPSP amplitude seen upon electrical stimulation of the perforant path in the ventral dentate gyrus in wildtype C57Bl6/J mice. Note the increase in amplitude upon superfusion of the mast cell degranulator C48/80 (1 μg/ml) and subsequent decrease in amplitude after the addition of histamine (10 μM) to the recording aCSF (grey trace). The addition of the H_3_ receptor antagonist thioperamide (10 μM) (blue trace) does not block the C48/80-mediated increase in amplitude but reduces the subsequent histamine-mediated decrease in amplitude. (**B**) Repeating the experiment in brain sections from Kit<W-sh> transgenic mice, which are largely deficient for brain mast cells, does not generate a C48/80-mediated increase in the fEPSP amplitude, but additional histamine (10 μM) superfusion led to a trend towards a decrease in their amplitude. (**C**) Graph of the normalised fEPSP amplitude upon stimulation of the more distal CA1 Schaffer collateral pathway and superfusion of C48/80 alone or in combination with histamine. Note the absence of an increase in fEPSP amplitude with C48/80 alone but a trend towards a reduction in amplitude upon washin of histamine. All data normalized to stable baseline and presented as mean ± SEM. ns, not significant; ** p<0.01

## Discussion

Here we demonstrate the presence of a transient and relatively large population of mast cells at the hippocampal-thalamic border as compared to other brain regions during the first postnatal weeks in C57Bl6/J mice. Moreover, we show that these mast cells when degranulated using the mast cell degranulator C48/80 results in an increase in synaptic transmission at the perforant pathway to dentate granule cell synapses. This might be mediated independently of histamine as the H_3_ receptor antagonist thioperamide does not block this increase in postsynaptic response with histamine superfusion tending to lead to a reduction in synaptic transmission which is sensitive to thioperamide.

It has been well described that mast cells populate the CNS and that their numbers fluctuate as a result of physiological, developmental and pathological changes and that they can impact neurons and other resident cells (Silver & Curley, 2013; Forsythe, 2019). Many of the compounds released by mast cells act in a paracrine fashion on these cells and therefore their localisation in the brain is important to comprehend (Forsythe, 2019). Here the presence of mast cells was studied in the brains of C57Bl/6 mice from the last week of embryogenesis until young adulthood. An initial focus was on striatum which we previously observed does not contain dense innervation of histaminergic afferents at young ages, but during this period does express histamine receptors and exhibits significant histaminergic modulation of neurons and synapses (Han *et al*., 2020; Márquez-Gómez *et al*., 2023). This raises the question what other sources of histamine might exists that impact this brain region, and we explored whether mast cells could contribute. We find that the largest number of CNS mast cells in C57Bl/6 mice are found during the first postnatal week and specifically between postnatal day (P)3-6 and below the hippocampus and that other brain regions including the striatum have significantly lower numbers. Previous studies also investigated mast cell localisation in the developing brain, but were often performed in rats, using various approaches including expression of the histamine synthesizing enzyme histidine decarboxylase (Karlstedt *et al*., 2001), histamine immunolabelling (Auvinen & Panula, 1988; Panula *et al*., 1988), toluidine blue (Ferrer *et al*., 1979; Lambracht-Hall *et al*., 1990a) and/or other staining (Hendrix *et al*., 2006; Khalil *et al*., 2007; Lenz *et al*., 2018; Blanchard *et al*., 2025). Our results using toluidine blue and avidin-FITC staining are in general agreement with these previous observations in that there is a transient period when mast cells located below the hippocampus are particularly prominent. Although others have found in rats and ICR albino mice (Yang *et al*., 1999) that they afterwards appear to settle in thalamic regions (Auvinen & Panula, 1988; Panula *et al*., 1988; Lambracht-Hall *et al*., 1990a; Khalil *et al*., 2007) likely through migration from pial regions towards the dorsal thalamic perivascular space and possibly along large blood vessels (Lambracht-Hall *et al*., 1990a), we did not observe this clearly in C57Bl6/J mice. However, we cannot exclude the possibility that methodological differences such as the rapid decapitation and immersion fixation of mouse brains here might have led to premature degranulation of mast cells which could account for lower numbers observed. Indeed, future experiments could incorporate mast cell stabilisers such as disodium cromoglycate (Dong *et al*., 2017) during slicing. To further test our observations, we also made use of a transgenic mouse line containing the *Kit^W-sh^* (or “sash”) mutation which results in an embryonic deficit and significant reduced numbers of mast cells soon after birth. Indeed, in these transgenic mice we did not observe mast cells as revealed with avidin-FITC in the CNS regions studied.

We next explored whether these peri-hippocampal mast cells could have a physiological impact on developing synapses after their degranulation using the classic mast cell secretagogue C48/80 (Paton, 1951). Numerous studies using C48/80 in tissue or animal models have contributed to a general view that observed effects are through mast cell degranulation. However, only a few studies explicitly confirm this mode of action in isolated mast cells or explore whether effects are reversed by mast cell stabilizers (Takeuchi *et al*., 1985; Chiang *et al*., 2000; Theoharides *et al*., 2000). Indeed, although there is evidence that it leads to brain mast cell degranulation (Ibrahim *et al*., 1980; Dimitriadou *et al*., 1990; Lambracht-Hall *et al*., 1990b), while not impacting microglia or pial macrophages (Ibrahim *et al*., 1980; Dimitriadou *et al*., 1990), there is also evidence that C48/80 can directly impact nervous transmission in enteric nerves and visceral afferents via a mechanism that is independent of mast cells (Schemann *et al*., 2012). Indeed, C48/80 has a propensity to elicit off-target effects depending on the concentration used and can exert other cellular effects including directly affecting neurotransmitter release (Lovenberg & Cubeddu, 1988; Heppner & Fiekers, 1992; Palomaki & Laitinen, 2006; Byrne *et al*., 2007; Schemann *et al*., 2012). We find that superfusion of a relatively low concentration (0.1 μg/ml) of C48/80 leads to an increase in the fEPSP amplitude at the perforant path to the dentate gyrus synapses and we do not observe this effect in slices made from *Kit^W-sh^* transgenic mice which are largely deficient for mast cells which would support the conclusion that observations are likely mast cell mediated. However, concerns have been raised regarding the specificity of the *Kit^W-sh^* (or “sash”) transgenic mouse line as a model of mast cell deficiency in that development of other cell types besides mast cells might also be disrupted by altered stem cell factor – c-kit receptor interactions (Nocka *et al*., 1989; Manova & Bachvarova, 1991; Zhang & Fedoroff, 1997; Galli *et al*., 2015), including neurons and neuronal progenitors (Nautiyal *et al*., 2012; Wasielewska *et al*., 2017). Interestingly, mast cell-deficient mice have marked deficits in hippocampal volume and cell proliferation, as well as altered hippocampus-dependent behaviours that can be reversed by increasing mast cell degranulation (Nautiyal *et al*., 2012). Therefore, we cannot currently exclude the possibility that neurons and/or synapses of the perforant path might be altered in the *Kit^W-sh^* transgenic mouse line as compared to C57Bl/6 and might respond differently to C48/80 and this should be further explored.

Our observed effects of mast cell degranulation may not be histaminergic. Indeed, we find that when C48/80 is superfused with the H_3_ receptor antagonist thioperamide this does not block the observed increase in fEPSP amplitude while reducing the negative modulation of synaptic transmission resulting from histamine superfusion. Although preliminary, these results would suggest that the effect might not be mediated through H_3_ receptors but instead through other histamine receptor subtypes or different released mediators. Mast cells release a host of chemical mediators, each with distinct signalling properties, and some with the capacity to alter the activity of neuronal circuits. For example, when mast cell-deficient mice exhibited deficits in hippocampal-associated behaviours and neurogenesis, this could be reversed by the chronic administration of a selective serotonin reuptake inhibitor, demonstrating that mast cell-derived serotonin is accountable for behaviourally altering hippocampal circuitry (Nautiyal *et al*., 2012). However, our observed increase in fEPSP amplitude reported here would not be consistent with modulation via the neuromodulators dopamine or serotonin as this most likely would lead to a decrease in synaptic transmission (Otmakhova & Lisman, 1999; Nozaki *et al*., 2016).

In conclusion, these findings highlight the potential for the degranulation of brain-resident mast cells to modulate developing synapses. Degranulation and local release of mediators could occur upon inflammation or triggered by acute stress (Theoharides *et al*., 1995; Baldwin, 2006). Given their role in neuroinflammation and numerous cellular interactions it has been postulated that brain mast cells could also be implicated in neurodevelopmental disorders (Theoharides *et al*., 2013; Song *et al*., 2020) but their role in this regard if any remains unclear. While histamine is among the mediators released, our data so far would suggest that the observed effects on hippocampal synaptic transmission are unlikely to be mediated via the H_3_ receptor. Thus, although mast cell degranulation has been shown to significantly contribute to overall CNS histamine levels, direct synaptic effects of mast cell-derived histamine remain to be determined.

## Acknowledgements

This work is supported through an MRC Career Development Award (MR/M009599/1) and a John Fell Fund award (162/059) to TE. Erasmus scholarship studentship to HGA. We are grateful for the advice on antibodies and supplied tissue from a *Mrgprb2-cre:tdTomato* transgenic mouse by Samuel Van Remoortel and Jean-Pierre Timmermans and help by Ricardo Marquez-Gomez.

## References

Abraham, D., Oster, H., Huber, M. & Leitges, M. (2007) The expression pattern of three mast cell specific proteases during mouse development. Mol Immunol, 44, 732–740.

Arias-Montano, J.A., Floran, B., Garcia, M., Aceves, J. & Young, J.M. (2001) Histamine H(3) receptor-mediated inhibition of depolarization-induced, dopamine D(1) receptor-dependent release of [(3)H]-gamma-aminobutryic acid from rat striatal slices. British journal of pharmacology, 133, 165–171.

Arrang, J.M., Drutel, G. & Schwartz, J.C. (1995) Characterization of histamine H3 receptors regulating acetylcholine release in rat entorhinal cortex. British journal of pharmacology, 114, 1518–1522.

Arrang, J.M., Garbarg, M., Lancelot, J.C., Lecomte, J.M., Pollard, H., Robba, M., Schunack, W. & Schwartz, J.C. (1987) Highly potent and selective ligands for histamine H3-receptors. Nature, 327, 117–123.

Auvinen, S. & Panula, P. (1988) Development of histamine-immunoreactive neurons in the rat brain. The Journal of comparative neurology, 276, 289–303.

Baldan, L.C., Williams, K.A., Gallezot, J.D., Pogorelov, V., Rapanelli, M., Crowley, M., Anderson, G.M., Loring, E., Gorczyca, R., Billingslea, E., Wasylink, S., Panza, K.E., Ercan-Sencicek, A.G., Krusong, K., Leventhal, B.L., Ohtsu, H., Bloch, M.H., Hughes, Z.A., Krystal, J.H., Mayes, L., de Araujo, I., Ding, Y.S., State, M.W. & Pittenger, C. (2014) Histidine decarboxylase deficiency causes tourette syndrome: parallel findings in humans and mice. Neuron, 81, 77–90.

Baldwin, A.L. (2006) Mast cell activation by stress. Methods Mol Biol, 315, 349–360.

Bekkers, J.M. (1993) Enhancement by histamine of NMDA-mediated synaptic transmission in the hippocampus. Science, 261, 104–106.

Bergstresser, P.R., Tigelaar, R.E. & Tharp, M.D. (1984) Conjugated avidin identifies cutaneous rodent and human mast cells. J Invest Dermatol, 83, 214–218.

Blanchard, A.C., Maximova, A., Phillips-Jones, T., Bruce, M.R., Anastasiadis, P., Dionisos, C.V., Engel, K., Reinl, E., Pham, A., Malaiya, S., Singh, N., Ament, S. & McCarthy, M.M. (2025) Mast cells proliferate in the peri-hippocampal space during early development and modulate local and peripheral immune cells. Dev Cell, 60, 853–870 e857.

Brown, R.E., Fedorov, N.B., Haas, H.L. & Reymann, K.G. (1995) Histaminergic modulation of synaptic plasticity in area CA1 of rat hippocampal slices. Neuropharmacology, 34, 181–190.

Byrne, R.D., Rosivatz, E., Parsons, M., Larijani, B., Parker, P.J., Ng, T. & Woscholski, R. (2007) Differential activation of the PI 3-kinase effectors AKT/PKB and p70 S6 kinase by compound 48/80 is mediated by PKCalpha. Cell Signal, 19, 321–329.

Carthy, E. & Ellender, T. (2021) Histamine, Neuroinflammation and Neurodevelopment: A Review. Frontiers in neuroscience, 15, 680214.

Chiang, G., Patra, P., Letourneau, R., Jeudy, S., Boucher, W., Green, M., Sant, G.R. & Theoharides, T.C. (2000) Pentosanpolysulfate inhibits mast cell histamine secretion and intracellular calcium ion levels: an alternative explanation of its beneficial effect in interstitial cystitis. J Urol, 164, 2119–2125.

Dimitriadou, V., Lambracht-Hall, M., Reichler, J. & Theoharides, T.C. (1990) Histochemical and ultrastructural characteristics of rat brain perivascular mast cells stimulated with compound 48/80 and carbachol. Neuroscience, 39, 209–224.

Dong, H., Zhang, X., Wang, Y., Zhou, X., Qian, Y. & Zhang, S. (2017) Suppression of Brain Mast Cells Degranulation Inhibits Microglial Activation and Central Nervous System Inflammation. Mol Neurobiol, 54, 997–1007.

Ellender, T.J., Huerta-Ocampo, I., Deisseroth, K., Capogna, M. & Bolam, J.P. (2011) Differential modulation of excitatory and inhibitory striatal synaptic transmission by histamine. J Neurosci, 31, 15340–15351.

Ercan-Sencicek, A.G.S., A.A.; Ghosh, A.K.; Bilguvar, K.’ O’Roak, B.J.; Mason, C.E.; Abbott, T.; Gupta, A.; King, R.A.; Pauls, D.L.; Tischfield, J.A.; Heiman, G.A.; Singer, H.S., Gilbert, D.L.; Hoekstra, P.J.; Morgan, T.M., Loring, E.; Yasuno, K.; Fernandez, T.; Sanders, S.; Louvi, A.; Cho, J.H.; Mane, S.; Colangelo, C.M.; Biederer, T.; Lifton, R.P.; Gunel, M.; State, M.W. (2010) L-histidine decarboxylase and tourettes syndrome. The New England journal of medicine, 363, 1901–1908.

Fernandez, T.V., Sanders, S.J., Yurkiewicz, I.R., Ercan-Sencicek, A.G., Kim, Y.S., Fishman, D.O., Raubeson, M.J., Song, Y., Yasuno, K., Ho, W.S., Bilguvar, K., Glessner, J., Chu, S.H., Leckman, J.F., King, R.A., Gilbert, D.L., Heiman, G.A., Tischfield, J.A., Hoekstra, P.J., Devlin, B., Hakonarson, H., Mane, S.M., Gunel, M. & State, M.W. (2012) Rare copy number variants in tourette syndrome disrupt genes in histaminergic pathways and overlap with autism. Biol Psychiatry, 71, 392–402.

Ferrer, I., Picatoste, F., Rodergas, E., Garcia, A., Sabria, J. & Blanco, I. (1979) Histamine and mast cells in developing rat brain. Journal of neurochemistry, 32, 587–592.

Flores, J.A., Ramirez-Ponce, M.P., Montes, M.A., Balseiro-Gomez, S., Acosta, J., Alvarez de Toledo, G. & Ales, E. (2019) Proteoglycans involved in bidirectional communication between mast cells and hippocampal neurons. J Neuroinflammation, 16, 107.

Forsythe, P. (2019) Mast Cells in Neuroimmune Interactions. Trends in neurosciences, 42, 43–55.

Franklin, K.B.J., Paxinos, G. (2007) The mouse brain atlas in stereotaxic coordinates. *Elsevier*.

Galli, S.J., Tsai, M., Marichal, T., Tchougounova, E., Reber, L.L. & Pejler, G. (2015) Approaches for analyzing the roles of mast cells and their proteases in vivo. Adv Immunol, 126, 45–127.

Gbahou, F., Vincent, L., Humbert-Claude, M., Tardivel-Lacombe, J., Chabret, C. & Arrang, J.M. (2006) Compared pharmacology of human histamine H3 and H4 receptors: structure-activity relationships of histamine derivatives. British journal of pharmacology, 147, 744–754.

Grimbaldeston, M.A., Chen, C.C., Piliponsky, A.M., Tsai, M., Tam, S.Y. & Galli, S.J. (2005) Mast cell-deficient W-sash c-kit mutant Kit W-sh/W-sh mice as a model for investigating mast cell biology in vivo. Am J Pathol, 167, 835–848.

Haas, H. & Panula, P. (2003) The role of histamine and the tuberomamillary nucleus in the nervous system. Nature reviews. Neuroscience, 4, 121–130.

Haas, H.L., Sergeeva, O.A. & Selbach, O. (2008) Histamine in the nervous system. Physiol Rev, 88, 1183–1241.

Han, S., Marquez-Gomez, R., Woodman, M. & Ellender, T. (2020) Histaminergic Control of Corticostriatal Synaptic Plasticity during Early Postnatal Development. J Neurosci, 40, 6557–6571.

Hendrix, S., Warnke, K., Siebenhaar, F., Peters, E.M., Nitsch, R. & Maurer, M. (2006) The majority of brain mast cells in B10.PL mice is present in the hippocampal formation. Neuroscience letters, 392, 174–177.

Heppner, T.J. & Fiekers, J.F. (1992) Compound 48/80 blocks transmission and increases the excitability of ganglion neurons. Eur J Pharmacol, 213, 427–434.

Hew, R.W., Hodgkinson, C.R. & Hill, S.J. (1990) Characterization of histamine H3-receptors in guinea-pig ileum with H3-selective ligands. British journal of pharmacology, 101, 621–624.

Ibrahim, M.Z., Uthman, M.A., Tenekjian, V. & Wiedman, T. (1980) The mast cells of the mammalian central nervous system. V. The effect of compound 48/80 on the neurolipomastocytoid cells and related areas of the CNS: early changes. Cell Tissue Res, 212, 99–116.

Joulia, R., Gaudenzio, N., Rodrigues, M., Lopez, J., Blanchard, N., Valitutti, S. & Espinosa, E. (2015) Mast cells form antibody-dependent degranulatory synapse for dedicated secretion and defence. Nature communications, 6, 6174.

Kafitz, K.W., Meier, S.D., Stephan, J. & Rose, C.R. (2008) Developmental profile and properties of sulforhodamine 101--Labeled glial cells in acute brain slices of rat hippocampus. Journal of neuroscience methods, 169, 84–92.

Karagiannidis, I., Dehning, S., Sandor, P., Tarnok, Z., Rizzo, R., Wolanczyk, T., Madruga-Garrido, M., Hebebrand, J., Nothen, M.M., Lehmkuhl, G., Farkas, L., Nagy, P., Szymanska, U., Anastasiou, Z., Stathias, V., Androutsos, C., Tsironi, V., Koumoula, A., Barta, C., Zill, P., Mir, P., Muller, N., Barr, C. & Paschou, P. (2013) Support of the histaminergic hypothesis in Tourette syndrome: association of the histamine decarboxylase gene in a large sample of families. J Med Genet, 50, 760–764.

Karlstedt, K., Nissinen, M., Michelsen, K.A. & Panula, P. (2001) Multiple sites of L-histidine decarboxylase expression in mouse suggest novel developmental functions for histamine. Dev Dyn, 221, 81–91.

Khalil, M., Ronda, J., Weintraub, M., Jain, K., Silver, R. & Silverman, A.-J. (2007) Brain mast cell relationship to neurovasculature during development. Brain research, 1171, 18–29.

Kierdorf, K., Masuda, T., Jordao, M.J.C. & Prinz, M. (2019) Macrophages at CNS interfaces: ontogeny and function in health and disease. Nature reviews. Neuroscience, 20, 547–562.

Kitamura, Y., Go, S. & Hatanaka, K. (1978) Decrease of mast cells in W/Wv mice and their increase by bone marrow transplantation. Blood, 52, 447–452.

Kovacs, P., Hernadi, I. & Wilhelm, M. (2006) Mast cells modulate maintained neuronal activity in the thalamus in vivo. J Neuroimmunol, 171, 1–7.

Lambracht-Hall, M., Dimitriadou, V. & Theoharides, T.C. (1990a) Migration of mast cells in the developing rat brain. Brain research. Developmental brain research, 56, 151–159.

Lambracht-Hall, M., Konstantinidou, A.D. & Theoharides, T.C. (1990b) Serotonin release from rat brain mast cells in vitro. Neuroscience, 39, 199–207.

Lenz, K.M., Pickett, L.A., Wright, C.L., Davis, K.T., Joshi, A. & McCarthy, M.M. (2018) Mast Cells in the Developing Brain Determine Adult Sexual Behavior. The Journal of neuroscience : the official journal of the Society for Neuroscience, 38, 8044–8059.

Lin, W., Xu, L., Zheng, Y., An, S., Zhao, M., Hu, W., Li, M., Dong, H., Li, A., Li, Y., Gong, H., Pan, G., Wang, Y., Luo, Q. & Chen, Z. (2023) Whole-brain mapping of histaminergic projections in mouse brain. Proceedings of the National Academy of Sciences of the United States of America, 120, e2216231120.

Lovenberg, T.W. & Cubeddu, L.X. (1988) Monoamine release by compound 48/80 from nonmast cell compartments in mouse brain slices. The Journal of pharmacology and experimental therapeutics, 247, 562–568.

Lucaci, D., Yu, X., Chadderton, P., Wisden, W. & Brickley, S.G. (2023) Histamine Release in the Prefrontal Cortex Excites Fast-Spiking Interneurons while GABA Released from the Same Axons Inhibits Pyramidal Cells. The Journal of neuroscience : the official journal of the Society for Neuroscience, 43, 187–198.

Lyon, M.F. & Glenister, P.H. (1982) A new allele sash (Wsh) at the W-locus and a spontaneous recessive lethal in mice. Genet Res, 39, 315–322.

Ma, Q., Jiang, L., Chen, H., An, D., Ping, Y., Wang, Y., Dai, H., Zhang, X., Wang, Y., Chen, Z. & Hu, W. (2023) Histamine H(2) receptor deficit in glutamatergic neurons contributes to the pathogenesis of schizophrenia. Proceedings of the National Academy of Sciences of the United States of America, 120, e2207003120.

Manova, K. & Bachvarova, R.F. (1991) Expression of c-kit encoded at the W locus of mice in developing embryonic germ cells and presumptive melanoblasts. Dev Biol, 146, 312–324.

Márquez-Gómez, R., Parke, B., Cras, Y., Gullino, S.L., Hashemi, P. & Ellender, T. (2023) Histamine originating from the BNST modulates corticostriatal synaptic transmission during early postnatal development. bioRxiv, 2023.2010.2019.563087.

McNeil, B.D., Pundir, P., Meeker, S., Han, L., Undem, B.J., Kulka, M. & Dong, X. (2015) Identification of a mast-cell-specific receptor crucial for pseudo-allergic drug reactions. Nature, 519, 237–241.

Molina-Hernández, A., Díaz, N.F. & Arias-Montaño, J.A. (2012) Histamine in brain development. J Neurochem, 122, 872–882.

Molina-Hernandez, A., Nunez, A., Sierra, J.J. & Arias-Montano, J.A. (2001) Histamine H3 receptor activation inhibits glutamate release from rat striatal synaptosomes. Neuropharmacology, 41, 928–934.

Molina-Hernández, A. & Velasco, I. (2008) Histamine induces neural stem cell proliferation and neuronal differentiation by activation of distinct histamine receptors. Journal of neurochemistry, 106, 706–717.

Morisset, S., Rouleau, A., Ligneau, X., Gbahou, F., Tardivel-Lacombe, J., Stark, H., Schunack, W., Ganellin, C.R., Schwartz, J.C. & Arrang, J.M. (2000) High constitutive activity of native H3 receptors regulates histamine neurons in brain. Nature, 408, 860–864.

Mukai, K., Chinthrajah, R.S., Nadeau, K.C., Tsai, M., Gaudenzio, N. & Galli, S.J. (2017) A new fluorescent-avidin-based method for quantifying basophil activation in whole blood. J Allergy Clin Immunol, 140, 1202–1206 e1203.

Nautiyal, K.M., Dailey, C.A., Jahn, J.L., Rodriquez, E., Son, N.H., Sweedler, J.V. & Silver, R. (2012) Serotonin of mast cell origin contributes to hippocampal function. The European journal of neuroscience, 36, 2347–2359.

Nocka, K., Majumder, S., Chabot, B., Ray, P., Cervone, M., Bernstein, A. & Besmer, P. (1989) Expression of c-kit gene products in known cellular targets of W mutations in normal and W mutant mice--evidence for an impaired c-kit kinase in mutant mice. Genes Dev, 3, 816–826.

Nozaki, K., Kubo, R. & Furukawa, Y. (2016) Serotonin modulates the excitatory synaptic transmission in the dentate granule cells. Journal of neurophysiology, 115, 2997–3007.

Otmakhova, N.A. & Lisman, J.E. (1999) Dopamine selectively inhibits the direct cortical pathway to the CA1 hippocampal region. The Journal of neuroscience : the official journal of the Society for Neuroscience, 19, 1437–1445.

Palomaki, V.A. & Laitinen, J.T. (2006) The basic secretagogue compound 48/80 activates G proteins indirectly via stimulation of phospholipase D-lysophosphatidic acid receptor axis and 5-HT1A receptors in rat brain sections. British journal of pharmacology, 147, 596–606.

Panula, P., Häppölä, O., Airaksinen, M.S., Auvinen, S. & Virkamäki, A. (1988) Carbodiimide as a tissue fixative in histamine immunohistochemistry and its application in developmental neurobiology. J Histochem Cytochem, 36, 259–269.

Panula, P., Sundvik, M. & Karlstedt, K. (2014) Developmental roles of brain histamine. Trends Neurosci, 37, 159–168.

Paton, W.D. (1951) Compound 48/80: a potent histamine liberator. Br J Pharmacol Chemother, 6, 499–508.

Quintana, F.J. (2019) Myeloid cells in the central nervous system: So similar, yet so different. Sci Immunol, 4.

Rapanelli, M., Frick, L., Bito, H. & Pittenger, C. (2017) Histamine modulation of the basal ganglia circuitry in the development of pathological grooming. Proceedings of the National Academy of Sciences of the United States of America, 114, 6599–6604.

Rapanelli, M. & Pittenger, C. (2016) Histamine and histamine receptors in Tourette syndrome and other neuropsychiatric conditions. Neuropharmacology, 106, 85–90.

Reiner, P.B.S., K.; Fibiger, H.C.; McGeer, E.G. (1988) Ontogeny of histidine-decarboxylase-immunoreactive neurons in the tuberomammillary nucleus of the rat hypothalamus: Time of origin and Development of transmitter phenotype. The Journal of comparative neurology, 276, 304–311.

Robertson, M.M., Eapen, V., Singer, H.S., Martino, D., Scharf, J.M., Paschou, P., Roessner, V., Woods, D.W., Hariz, M., Mathews, C.A., Crncec, R. & Leckman, J.F. (2017) Gilles de la Tourette syndrome. Nat Rev Dis Primers, 3, 16097.

Schemann, M., Kugler, E.M., Buhner, S., Eastwood, C., Donovan, J., Jiang, W. & Grundy, D. (2012) The mast cell degranulator compound 48/80 directly activates neurons. PLoS One, 7, e52104.

Schwartz, J.C., Arrang, J.M., Garbarg, M., Pollard, H. & Ruat, M. (1991) Histaminergic transmission in the mammalian brain. Physiol Rev, 71, 1–51.

Shan, L., Bao, A.M. & Swaab, D.F. (2015) The human histaminergic system in neuropsychiatric disorders. Trends in neurosciences, 38, 167–177.

Silver, R. & Curley, J.P. (2013) Mast cells on the mind: new insights and opportunities. Trends in neurosciences, 36, 513–521.

Silver, R., Silverman, A.J., Vitkovic, L. & Lederhendler, II (1996) Mast cells in the brain: evidence and functional significance. Trends in neurosciences, 19, 25–31.

Song, Y., Lu, M., Yuan, H., Chen, T. & Han, X. (2020) Mast cell-mediated neuroinflammation may have a role in attention deficit hyperactivity disorder (Review). Exp Ther Med, 20, 714–726.

St John, A.L., Rathore, A.P.S. & Ginhoux, F. (2023) New perspectives on the origins and heterogeneity of mast cells. Nat Rev Immunol, 23, 55–68.

Takahashi, K., Lin, J.S. & Sakai, K. (2006) Neuronal activity of histaminergic tuberomammillary neurons during wake-sleep states in the mouse. The Journal of neuroscience : the official journal of the Society for Neuroscience, 26, 10292–10298.

Takeuchi, K., Ohtsuki, H., Nakagawa, S. & Okabe, S. (1985) Characterization of FPL-52694 [5-(2-hydroxypropoxyl)-8-propyl-4-oxo-4H-benzopyran-2-carboxylic acid Na] on histamine release from rat peritoneal mast cells induced by antigen, compound 48/80 and A 23187. Agents Actions, 17, 10–13.

Tharp, M.D., Seelig, L.L., Jr., Tigelaar, R.E. & Bergstresser, P.R. (1985) Conjugated avidin binds to mast cell granules. J Histochem Cytochem, 33, 27–32.

Theoharides, T.C., Asadi, S., Panagiotidou, S. & Weng, Z. (2013) The “missing link” in autoimmunity and autism: extracellular mitochondrial components secreted from activated live mast cells. Autoimmun Rev, 12, 1136–1142.

Theoharides, T.C., Patra, P., Boucher, W., Letourneau, R., Kempuraj, D., Chiang, G., Jeudy, S., Hesse, L. & Athanasiou, A. (2000) Chondroitin sulphate inhibits connective tissue mast cells. British journal of pharmacology, 131, 1039–1049.

Theoharides, T.C., Spanos, C., Pang, X., Alferes, L., Ligris, K., Letourneau, R., Rozniecki, J.J., Webster, E. & Chrousos, G.P. (1995) Stress-induced intracranial mast cell degranulation: a corticotropin-releasing hormone-mediated effect. Endocrinology, 136, 5745–5750.

Theoharides, T.C., Twahir, A. & Kempuraj, D. (2024) Mast cells in the autonomic nervous system and potential role in disorders with dysautonomia and neuroinflammation. Ann Allergy Asthma Immunol, 132, 440–454.

Tsai, M., Valent, P. & Galli, S.J. (2022) KIT as a master regulator of the mast cell lineage. J Allergy Clin Immunol, 149, 1845–1854.

Valle-Bautista, R., Marquez-Valadez, B., Herrera-Lopez, G., Griego, E., Galvan, E.J., Diaz, N.F., Arias-Montano, J.A. & Molina-Hernandez, A. (2021) Long-Term Functional and Cytoarchitectonic Effects of the Systemic Administration of the Histamine H(1) Receptor Antagonist/Inverse Agonist Chlorpheniramine During Gestation in the Rat Offspring Primary Motor Cortex. Frontiers in neuroscience, 15, 740282.

Van Nassauw, L., Adriaensen, D. & Timmermans, J.P. (2007) The bidirectional communication between neurons and mast cells within the gastrointestinal tract. Auton Neurosci-Basic, 133, 91–103.

Vanhala, A., Yamatodani, A. & Panula, P. (1994) Distribution of histamine-, 5-hydroxytryptamine-, and tyrosine hydroxylase-immunoreactive neurons and nerve fibers in developing rat brain. The Journal of comparative neurology, 347, 101–114.

Wasielewska, J.M., Gronnert, L., Rund, N., Donix, L., Rust, R., Sykes, A.M., Hoppe, A., Roers, A., Kempermann, G. & Walker, T.L. (2017) Mast cells increase adult neural precursor proliferation and differentiation but this potential is not realized in vivo under physiological conditions. Sci Rep, 7, 17859.

Xu, L., Lin, W., Zheng, Y., Chen, J., Fang, Z., Tan, N., Hu, W., Guo, Y., Wang, Y. & Chen, Z. (2022) An H2R-dependent medial septum histaminergic circuit mediates feeding behavior. Current biology : CB, 32, 1937–1948 e1935.

Yamatodani, A., Maeyama, K., Watanabe, T., Wada, H. & Kitamura, Y. (1982) Tissue distribution of histamine in a mutant mouse deficient in mast cells: clear evidence for the presence of non-mast-cell histamine. Biochem Pharmacol, 31, 305–309.

Yamazaki, M., Tsujimura, T., Morii, E., Isozaki, K., Onoue, H., Nomura, S. & Kitamura, Y. (1994) C-kit gene is expressed by skin mast cells in embryos but not in puppies of Wsh/Wsh mice: age-dependent abolishment of c-kit gene expression. Blood, 83, 3509–3516.

Yang, M., Chien, C. & Lu, K. (1999) Morphological, immunohistochemical and quantitative studies of murine brain mast cells after mating. Brain research, 846, 30–39.

Zhang, S.C. & Fedoroff, S. (1997) Cellular localization of stem cell factor and c-kit receptor in the mouse nervous system. Journal of neuroscience research, 47, 1–15.

Zhang, X.Y., Peng, S.Y., Shen, L.P., Zhuang, Q.X., Li, B., Xie, S.T., Li, Q.X., Shi, M.R., Ma, T.Y., Zhang, Q., Wang, J.J. & Zhu, J.N. (2020) Targeting presynaptic H3 heteroreceptor in nucleus accumbens to improve anxiety and obsessive-compulsive-like behaviors. Proceedings of the National Academy of Sciences of the United States of America, 117, 32155–32164.

